# TMEM11-Dependent Cristae Architecture Restrains Imiquimod-Induced NLRP3 Inflammasome Activation via Mitochondrial ROS

**DOI:** 10.64898/2026.11.06.764965

**Authors:** Beibei Wu, Matthew Do, Cristiane Beninca, Vaithilingaraja Arumugaswami, Yousang Gwack, Sonal Srikanth

## Abstract

Imiquimod (IMQ) is a clinically approved drug widely used to treat skin malignancies; however, its application frequently provokes severe inflammation through NLRP3 inflammasome activation. The mechanism underlying IMQ-induced activation of the NLRP3 inflammasome is not completely defined. While IMQ-induced inhibition of mitochondrial electron transport chain complex I is necessary for activation of the NLRP3 inflammasome, it is not sufficient; suggesting presence of additional targets/triggers. The role of mitochondrial reactive oxygen species (mtROS) as the additional trigger to activate the inflammasome remains contentious based on conflicting reports. Using a unique genetic model lacking expression of mitochondrial membrane protein TMEM11 we reaffirm the role of mtROS in IMQ-induced NLRP3 inflammasome activation. Macrophages deficient in *Tmem11*, a mitochondrial contact site and cristae organizing system (MICOS)-associated membrane-protein, exhibit ultrastructural defects in cristae morphology without altering mitochondrial mass or mtDNA copy number. Loss of TMEM11 influences cristae length and density, reducing mitochondrial oxygen consumption. *Tmem11*-deficient macrophages exhibit normal NLRP3 inflammasome activation in response to canonical agonists (e.g., ATP, nigericin) but show caspase-1 hyperactivation and exaggerated IL-1β and IL-18 secretion upon IMQ treatment. This exaggerated response is rescued by antioxidant treatment, confirming the role for mtROS. These findings demonstrate that cristae architecture restrains drug-induced mtROS to limit NLRP3 activation and that mtROS is a primary driver of IMQ-mediated NLRP3 inflammasome activation.

## Introduction

The innate immune system relies on pattern recognition receptors to sense pathogen-associated molecular patterns and damage-associated molecular patterns. Among these cytosolic receptors, the NLRP3 inflammasome is unique in its ability to respond to a diverse array of crystalline agents, membrane-pore-forming toxins, and pharmacological compounds^1-3^. Upon activation, NLRP3 oligomerizes and recruits the adaptor protein apoptosis-associated speck-like protein containing a CARD (ASC) domain, which nucleates the assembly of pro-caspase-1 into active caspase-1. Active caspase-1 subsequently cleaves pro-IL-1β and pro-IL-18 into their mature, bioactive cytokine forms and processes gasdermin D to trigger pyroptotic cell death.

Imiquimod (IMQ), a synthetic small-molecule imidazoquinoline, is an FDA-approved topical immunomodulator widely used for the treatment of superficial basal cell carcinomas, actinic keratoses, and external genital warts^4^. Although its therapeutic efficacy relies on activating innate immunity via direct stimulation of TLR7^5^, excessive application or prolonged exposure to IMQ frequently provokes severe cutaneous erythema, scaling, epidermal thickening, and systemic psoriasis-like inflammatory disease^6^. IMQ drives this inflammation through both TLR7-dependent pathways and TLR7-independent mechanisms involving the direct inhibition of mitochondrial electron transport chain (ETC) complex I and quinone oxidoreductases^7, 8^. Mitochondrial perturbation caused by IMQ initiates rapid NLRP3 inflammasome assembly and downstream IL-1β and IL-18 secretion^8^. Despite widespread recognition that mitochondria act as pivotal signaling platforms for NLRP3 activation^9-11^, the exact upstream mitochondrial messengers governing IMQ-induced inflammasome assembly are not well-defined. Initial paradigms established that IMQ-induced inhibition of complex I triggers mitochondrial respiratory chain arrest, elevating mtROS generation to serve as the proximal trigger for NLRP3 assembly^8^. However, recent investigations have challenged this mechanism, proposing that inhibition of complex I in conjunction with cellular perturbation (e.g., endolysosomal stress), independent of mtROS, serves as the dominant signal driving IMQ-induced NLRP3 activation^12^.

A primary obstacle in resolving whether mtROS represents the definitive trigger is that existing genetic models of mitochondrial dysfunction lack structural precision. Widely studied genetic perturbations—such as ablation of optic atrophy 1 (*Opa1*), dynamin-related protein 1 (*Drp1*), or mitochondrial transcription factor A (*Tfam*)—disrupt outer or inner membrane dynamics, broadly deplete mitochondrial mass, degrade mtDNA integrity, and induce baseline bioenergetic collapse and apoptosis^13-15^. Consequently, it has remained impossible to determine the discrete contribution of inner mitochondrial membrane cristae architecture to inflammasome regulation under physiologically relevant conditions. Mitochondrial cristae are specialized invaginations of the inner mitochondrial membrane that serve as bioenergetic hubs housing respiratory chain complexes and ATP synthase dimers. The structural organization of cristae junctions and sheets is coordinated by the multiprotein mitochondrial contact site and cristae organizing system (MICOS) complex, together with OPA1 oligomers. TMEM11 is a mitochondrial membrane protein that physically and functionally associates with the MICOS complex^16, 17^. Our previous work has revealed that deletion of *Tmem11* disrupts cristae ultrastructure and substantially elevates mtROS production without compromising cell viability, proliferation, or total mitochondrial mass^18^.

In this study, we utilize primary bone marrow–derived macrophages lacking TMEM11 expression to interrogate how cristae architecture governs NLRP3 inflammasome activation. We demonstrate that TMEM11 deficiency impairs mitochondrial cristae density and respiratory capacity while maintaining normal mitochondrial mass and mtDNA content. While TMEM11 deficiency did not influence NLRP3 inflammasome activation in response to canonical agonists like ATP or Nigericin, treatment with IMQ caused selective hyperactivation of caspase-1 leading to excessive secretion of IL-1β and IL-18 driven by exaggerated mtROS production. Mitigating mtROS reversed this inflammatory hyperactivation. Our findings reaffirm the role of mtROS in drug-induced NLRP3 inflammasome activation and identify TMEM11-dependent cristae architecture as a distinct structural checkpoint that restrains drug-induced inflammatory pathology.

## Materials and Methods

### Mice and Primary Macrophage Culture

*Tmem11*^*−/−*^ mice in C57BL/6NCrl genetic background (TCPR0855_ADGQ) were purchased from The Center for Phenogenomics (ON, Canada). The animals were bred and genotyped as described recently^18^. 8-12-week-old age and sex matched control (*Tmem11*^*+/+*^) and *Tmem11*^*-/-*^ littermate animals were used for all primary macrophage preparations. Primary bone marrow–derived macrophages (BMDMs) were generated by flushing femoral and tibial marrow from control and *Tmem11*^*-/-*^ mice. Cells were cultured in RPMI supplemented with 10% heat-inactivated fetal bovine serum, 100 U/mL penicillin, 100 μg/mL streptomycin, 2 mM L-glutamine, 0.5 mM Na-pyruvate and 8-10% conditioned medium from recombinant CMG14-12 cell line expressing M-CSF (or 20 ng/mL recombinant murine M-CSF) for 6 days at 37°C in a humidified atmosphere of 5% CO2. All mice were maintained in pathogen-free barrier facilities and used in accordance with protocols approved by the Institutional Animal Care and Use Committee at the UCLA.

### Inflammasome Stimulation and Cytokine Quantification

Differentiated BMDMs were seeded into plates and primed with 100 ng/mL ultrapure lipopolysaccharide (LPS) from *Escherichia coli* O111:B4 for 3 h. Primed BMDMs were subsequently challenged with imiquimod (IMQ; 10–20 μg/mL) for 90 min, ATP (2–5 mM) for 30 min, or nigericin (5–15 μM) for 45 min. For assessment of non-canonical NLRP3 activation, LPS was delivered intracellularly by transfection using Lipofectamine 2000 (Thermo Fisher Scientific). For AIM2 inflammasome and TLR3 activation, cells were transfected with poly(dA:dT) or poly(I:C), respectively, complexed with Lipofectamine 2000 for 5 h. In antioxidant rescue experiments, LPS-primed cells were pre-treated with the indicated concentrations of N-acetylcysteine (NAC) for 30 min before IMQ stimulation. Cell-culture supernatants were harvested following stimulation and cleared of cellular debris by centrifugation for 5 min at 4°C. Secreted murine IL-1β and IL-18 were quantified using commercial sandwich ELISA kits (Thermo Fisher Scientific) according to the manufacturers’ instructions.

### Immunoblotting and mtDNA Copy Number Analysis

For analysis of caspase-1 cleavage, cell culture supernatants and cell lysates from IMQ-treated control and *Tmem11*^*-/-*^ BMDMs were combined. Supernatant proteins were concentrated, and whole-cell lysates were prepared in RIPA lysis buffer supplemented with protease and phosphatase inhibitors. Protein samples were resolved on SDS-PAGE gels, transferred onto 0.2 μm PVDF membranes, and probed with primary antibodies against caspase-1 (p20) (Cell Signaling Technology, clone E2G2I) and β-actin (Santa Cruz Biotechnology, #sc-47778). For analysis of mitochondrial proteins control and *Tmem11*^*-/-*^ BMDMs lysates were probed with antibodies targeting total OXPHOS rodent cocktail (CI-NDUFB8, CII-SDHB, CIII-UQCRC2, CIV-MTCO1, CV-ATP5A; Abcam, #MS604-300), TMEM11 (Proteintech, #16564-1-AP), MIC10 (Bioss antibodies, #BS-15029R), MIC60 (Proteintech, #10179-1-AP), OPA1 (BD biosciences, # 612606), TOMM20 (Abcam, #ab56783), and β-actin (Santa Cruz Biotechnology, #sc-47778). Membranes were washed, incubated with HRP-conjugated secondary antibodies, developed using ECL substrates, and analyzed via densitometry. Total genomic and mitochondrial DNA was extracted from control and *Tmem11*−/− BMDMs, and real-time quantitative PCR targeting three discrete non-overlapping regions of the mitochondrial genome was conducted to calculate relative mtDNA copy numbers normalized against nuclear gene loci, as previously described^18^.

### Confocal Microscopy, Electron Microscopy and Super-Resolution STED Nanoscopy

For confocal analysis, BMDMs were stained with 50 nM Mito-tracker green in Opti-MEM medium at 37°C for 30 mins, followed by Hoechst 33342 staining for 10 mins. Cells were kept in a humidified incubation chamber at 37°C with 5% CO2 during image collection. Samples were imaged on a Zeiss LSM 880 confocal microscope coupled with a 63x lens in Airyscan mode and processed using Zen Black software (Zeiss). For mitochondria morphology analysis, 3D confocal images were analyzed using the machine learning pixel classifier in Aivia v.10 software (Leica Microsystems). Labelled mitochondria (MitoTracker green) were used for segmentation of the mitochondrial network that was measured in the cellular 3D volume. For electron microscopy, BMDMs were fixed in 2.5% glutaraldehyde in 0.1 M sodium cacodylate buffer for 2 h, post-fixed in 1% osmium tetroxide containing 1.5% potassium ferrocyanide for 1 hour, dehydrated through graded ethanol series, infiltrated with epoxy resin, sectioned at 70 nm and mounted on 400 mesh copper grids with carbon support film. Micrographs were acquired using a mini transmission electron microscope (DeLong) operating at 25 kV. Morphometric parameters — including mitochondrial area, maximal cristae width, mean cristae length, and cristae density—were quantitatively evaluated using pixel classification training and segmentation in Aivia v.10 software across multiple biological replicates. For stimulated emission depletion (STED) nanoscopy, live primary macrophages were stained with 250 nM of Abberior Live Orange inner mitochondrial membrane fluorescent probe for 1 h in phenol red-free medium at 37°C (5% CO2), followed by a 20 min wash with medium. Cells were imaged live using the Abberior STEDycon system equipped with a 100×/1.45 N.A and a 775 nm pulsed STED laser. STED images were taken with a 30 nm pixel size and 10 ms dwell time. Analysis was performed in ImageJ by drawing a line in the middle of the mitochondria and counting peaks of fluorescence to determine the number of cristae per length of mitochondria imaged.

### Metabolic Profiling, ATP and mtROS Measurements

Mitochondrial respiration and glycolytic rates were determined using a Seahorse XFe96 Extracellular Flux Analyzer. Control and *Tmem11*^*-/-*^ BMDMs were primed with LPS (100 ng/mL, 3 h) and either left untreated or exposed to IMQ (10 μg/mL) for 1 h prior to sequential injection of oligomycin (1.5 μM), FCCP (1.0 μM), and rotenone/antimycin A (0.5 μM each). Basal oxygen consumption rate, ATP-linked respiration, maximal respiratory capacity, spare respiratory capacity, and extracellular acidification rates were calculated. Total intracellular ATP levels were measured in basal and LPS-primed states using a luciferin-luciferase bioluminescent assay kit (Promega). Mitochondrial superoxide production was measured by staining BMDMs with 5 μM MitoSOX Red for 15 min at 37°C, followed by flow-cytometric analysis using a BD LSRFortessa flow cytometer. Mean fluorescence intensity (MFI) was used as the measure of MitoSOX signal.

### Statistical Analysis

Data are presented as mean ± SEM of at least three independent biological replicates. Statistical comparisons between two experimental groups were performed using two-tailed unpaired Student’s *t*-tests. Multi-group comparisons were evaluated by one-way or two-way analysis of variance followed by Tukey’s or Sidak’s post hoc tests. Statistical significance was established at *P* < 0.05.

## Results and Discussion

### TMEM11 deficiency in macrophages remodels cristae architecture without influencing mitochondrial abundance

TMEM11-deficient T cells exhibit altered cristae architecture without gross mitochondrial abnormalities^18^. To delineate how mitochondrial cristae structure impacts innate immune activation, we evaluated primary BMDMs from *Tmem11*^*-/-*^ mice. Classical models of mitochondrial disruption (*Opa1, Drp1, Tfam* knockouts) generally trigger rapid mtDNA depletion, massive organellar fragmentation, and reduced cell viability. In contrast, *Tmem11* deletion yielded completely viable macrophages upon differentiation from bone marrow precursors. Confocal microscopy demonstrated similar numbers of total mitochondria between control and *Tmem11*−/− cells (**Fig. 1A**). However, morphometric assessment revealed structural remodeling, where mitochondrial length was significantly shortened, total mitochondrial area and perimeter per cell were substantially reduced, and mitochondrial circularity was elevated in *Tmem11*^-/-^ cells, reflecting a structural transition from elongated tubular network into discrete fragmented units.

**Figure 1.**
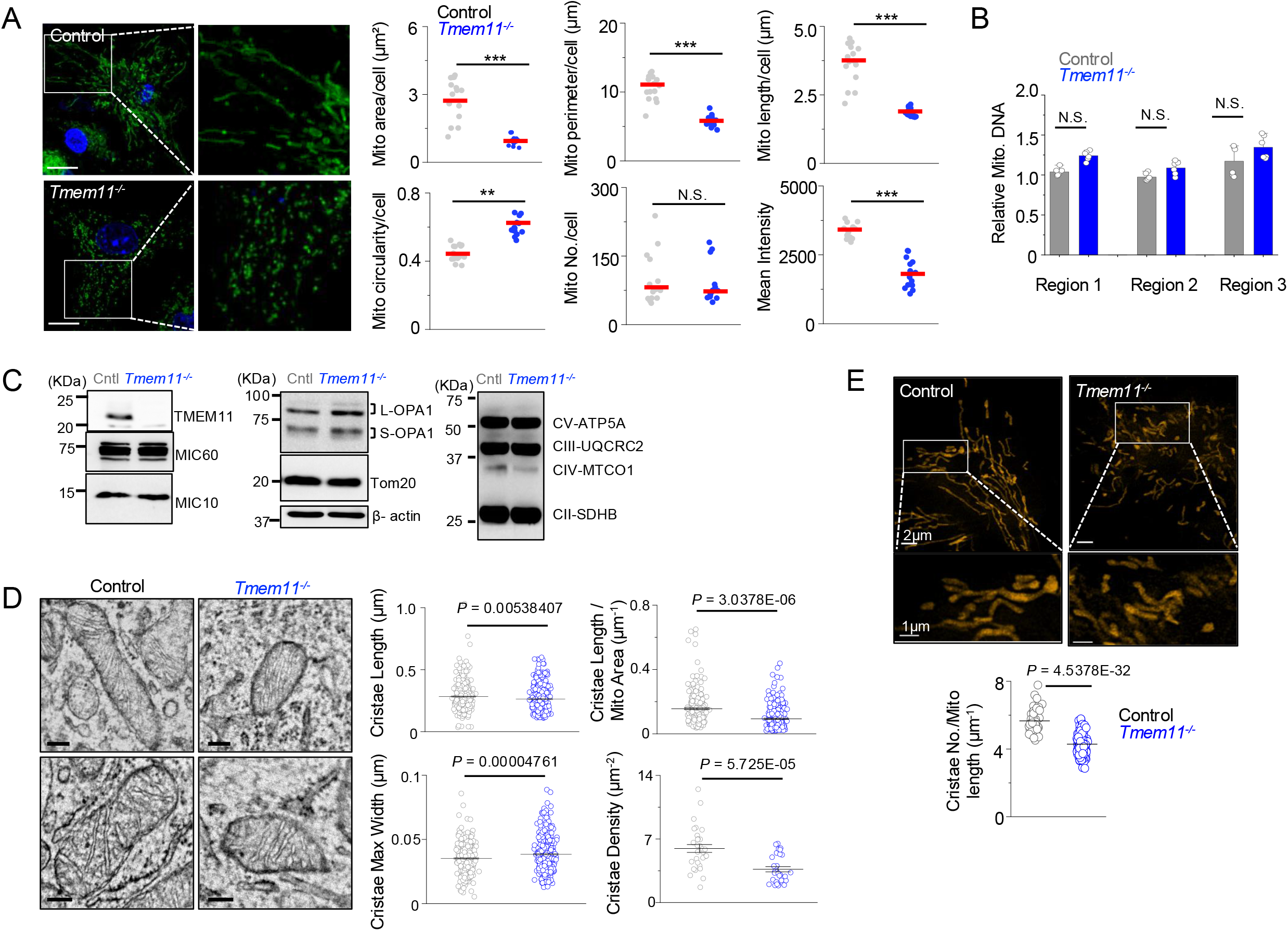
TMEM11 deficiency influences mitochondrial morphology and cristae architecture. **A**, Representative confocal images of WT and *Tmem11*^*-/-*^ primary BMDMs stained with MitoTracker green and Hoechst (nucleus). Scale bar, 2 µm. Scatterplots on the right show measurements of mitochondrial area, sphericity, length, and numbers. Each symbol represents data from an independent cell. **B**, Comparison of mitochondrial DNA content in WT and *Tmem11*^*-/-*^ BMDMs using quantitative PCR analysis of three different loci in the mitochondrial genome. Data represent triplicates from two independent experiments. **C**, Representative immunoblots of TMEM11, MIC10, MIC60, OPA1, TOMM20, β-actin, and mitochondrial respiratory-chain complex subunits (CII-SDHB, CIII-UQCRC2, CIV-MTCO1, and CV-ATP5A) from control and *Tmem11*^-/-^ BMDMs. **D**, Representative transmission electron micrographs of mitochondria from control and *Tmem11*^-/-^ BMDMs. Quantification of cristae length, maximal cristae width, cristae density, and cristae length normalized to mitochondrial area is shown. Scale bar, 100 nm. Each symbol represents data from an individual mitochondrion. **E**, Representative STED images of mitochondrial inner-membrane cristae in control and *Tmem11*^-/-^ BMDMs. Quantification of cristae number per mitochondrial length is shown. Scale bars, 2 μm and 1 μm as indicated. Each symbol represents data from an individual mitochondrion. All data are representative of three independent experiments (unless otherwise indicated) and graphs show means ± s.e.m.. *P < 0.05, **P < 0.005, ***P < 0.0005; N.S., not significant.

To confirm whether mitochondrial number was influenced, we evaluated relative mtDNA content. Quantitative PCR targeting three discrete non-overlapping regions of the mitochondrial genome revealed no significant differences between control and *Tmem11*^*-/-*^ macrophages (**Fig. 1B**). Immunoblot analysis confirmed absence of TMEM11 protein in knockout macrophages while demonstrating that levels of other cristae-shaping factors—including MICOS complex components MIC10 and MIC60, long and short OPA1 isoforms, and the outer mitochondrial membrane marker TOMM20—were similar to those in control cells (**Fig. 1C**). Immunoblot analysis also demonstrated that steady-state protein levels of the ETC complex subunits—including CII-SDHB, CIII-UQCRC2, CIV-MTCO1, and CV-ATP5A—were identical between control and *Tmem11*^*-/-*^ BMDMs.

Ultrastructure analysis using transmission electron microscopy revealed significant defects in mitochondrial inner membrane architecture of *Tmem11*^*-/-*^ BMDMs. While mitochondria from control macrophages possessed abundant, regular cristae spanning the matrix, those from *Tmem11*^-/-^ macrophages exhibited disorganized, sparse cristae sheets (**Fig. 1D**). Morphometric quantification showed that cristae density was reduced by ∼50%, total and normalized cristae length was decreased, and maximal cristae width was increased in mitochondria from *Tmem11*^*-/-*^ cells. Super-resolution STED nanoscopy verified that cristae number normalized to mitochondrial length was markedly reduced in *Tmem11*^*-/-*^ cells (**Fig. 1E**). These ultrastructural observations confirm that TMEM11 deficiency selectively impairs cristae invagination and architecture while preserving mitochondrial mass, genome copy numbers, as well as expression of key components of the MICOS and ETC complexes, and OPA1; establishing *Tmem11*^-/-^ primary BMDMs as a unique model to evaluate the role of cristae architecture in innate immune regulation.

### TMEM11 deficiency impairs respiration in macrophages

Cristae membranes serve as the physical scaffold for the spatial assembly and supercomplex stabilization of the mitochondrial electron transport chain. While expression of key components of the ETC complex was not altered in *Tmem11*^-/-^ BMDMs (**Fig. 1C**), their functional assembly may be influenced by altered cristae architecture. Therefore, we assessed whether cristae remodeling affected mitochondrial respiration and energy production. Seahorse metabolic flux profiling revealed that *Tmem11*^-/-^ BMDMs exhibited impaired mitochondrial oxygen consumption rates under basal conditions (**Fig. 2A**). Following sequential administration of oligomycin, FCCP, and rotenone/antimycin A, ATP-linked respiration was reduced, maximal uncoupled respiration dropped, and spare respiratory capacity showed a trend of reduction. Treatment with IMQ (10 μg/mL) further impaired the maximal respiration and spare respiratory capacity in *Tmem11*^-/-^ macrophages.

**Figure 2.**
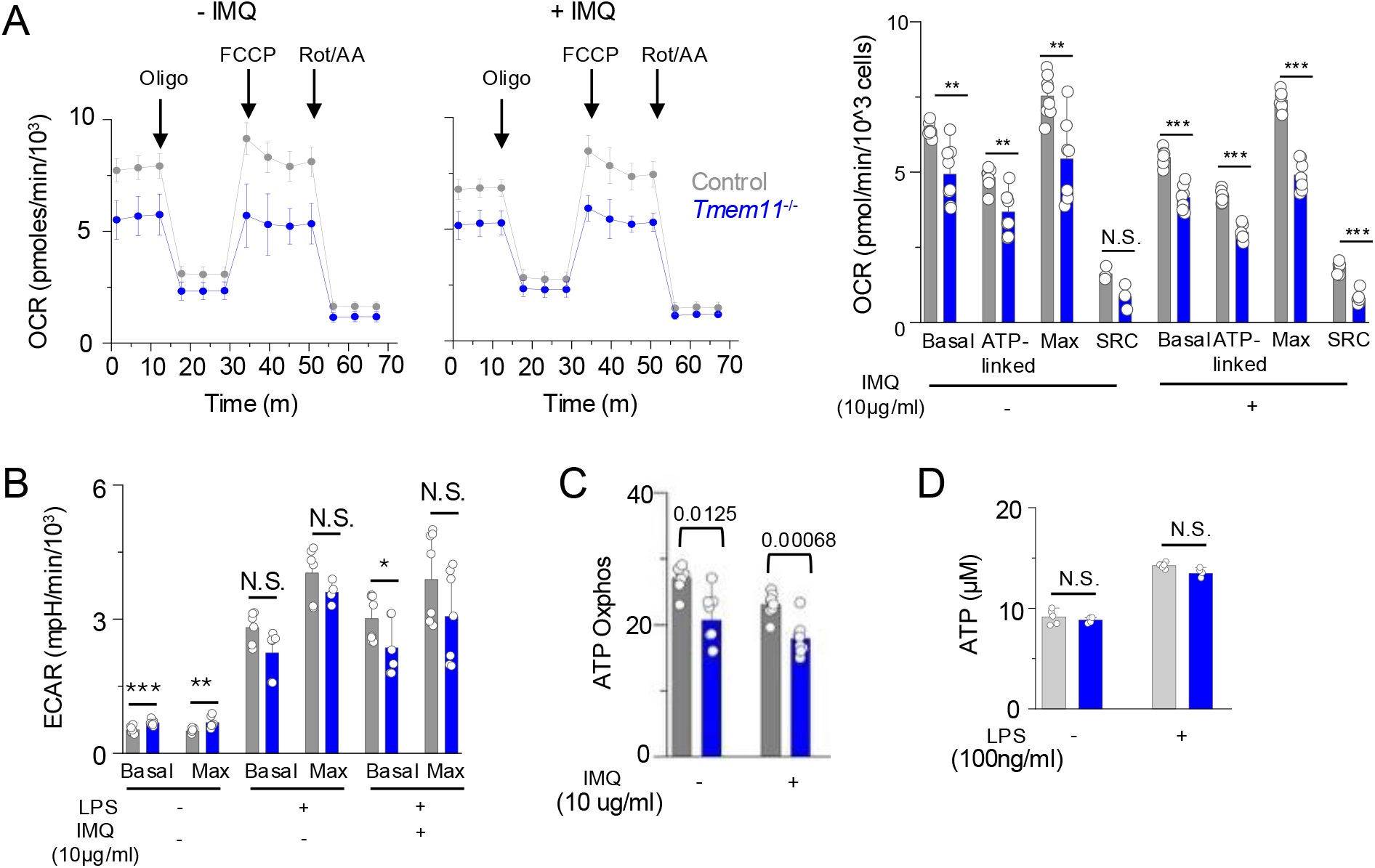
TMEM11 deficiency impairs mitochondrial respiration. **A**, Representative oxygen consumption rate (OCR) profiles of control and *Tmem11*^-/-^ BMDMs following sequential addition of oligomycin, FCCP, and rotenone/antimycin A (Rot/AA), in the absence or presence of IMQ (10 μg/mL). Bar graph on the right shows quantification of basal respiration, ATP-linked respiration, maximal respiration, and spare respiratory capacity (SRC). **B**, Bar graph shows baseline and maximal extracellular acidification rate (ECAR) in untreated, LPS and IMQ-treated control and *Tmem11*^-/-^ BMDMs. **C**, Seahorse analysis-derived ATP production rates attributable to oxidative phosphorylation (ATP Oxphos) in control and *Tmem11−/−* BMDMs under the indicated conditions. **D**, Total intracellular ATP concentrations measured by luciferase assay in control and *Tmem11−/−* BMDMs under basal and LPS-primed conditions. Data are presented as mean ± SEM from 2-3 independent biological replicates. Statistical significance was determined as described in the Methods. *P < 0.05, **P < 0.005, ***P < 0.0005; N.S., not significant.

Extracellular acidification rate measurements revealed that untreated *Tmem11*^-/-^ BMDMs mounted a compensatory increase in baseline and maximal glycolytic rates compared with control cells, indicating an adaptive bioenergetic shift toward glycolysis (**Fig. 2B**). However, this baseline glycolytic advantage was lost following LPS priming, which robustly accelerated glycolysis across control and *Tmem11*^-/-^ BMDMs to comparable levels. While real-time ATP generation rate analysis indicated a modest reduction in oxidative phosphorylation-derived ATP in *Tmem11*^-/-^ macrophages (**Fig. 2C**), total intracellular ATP levels remained preserved, showing no significant differences between genotypes in either the basal or LPS-primed state (**Fig. 2D**). These results suggest that altered cristae morphology in *Tmem11*^-/-^ macrophages hinders electron transport chain flux without reducing the abundance of respiratory complexes or depleting steady-state cellular ATP pools, likely due to compensatory increase in glycolytic flux.

### TMEM11 selectively restrains IMQ-induced NLRP3 inflammasome activation

Mitochondria are known to play a central role in various innate immune responses, including activation of the NLRP3 inflammasome^9^-^11, 17^. Hence, we tested the functional consequences of cristae disruption on innate immune activation by exposing LPS-primed BMDMs to various NLRP3 agonists. When challenged with canonical NLRP3 agonists, stimulation with extracellular ATP (2.5 and 5 mM) resulted in equivalent secretion of IL-1β and IL-18 in control and *Tmem11*^-/-^ macrophages (**Fig. 3A**). Another canonical NLRP3 agonist, the K^+^ ionophore nigericin (7.5 and 15 μM) also produced similar levels of IL-1β and IL-18 release across genotypes.

**Figure 3.**
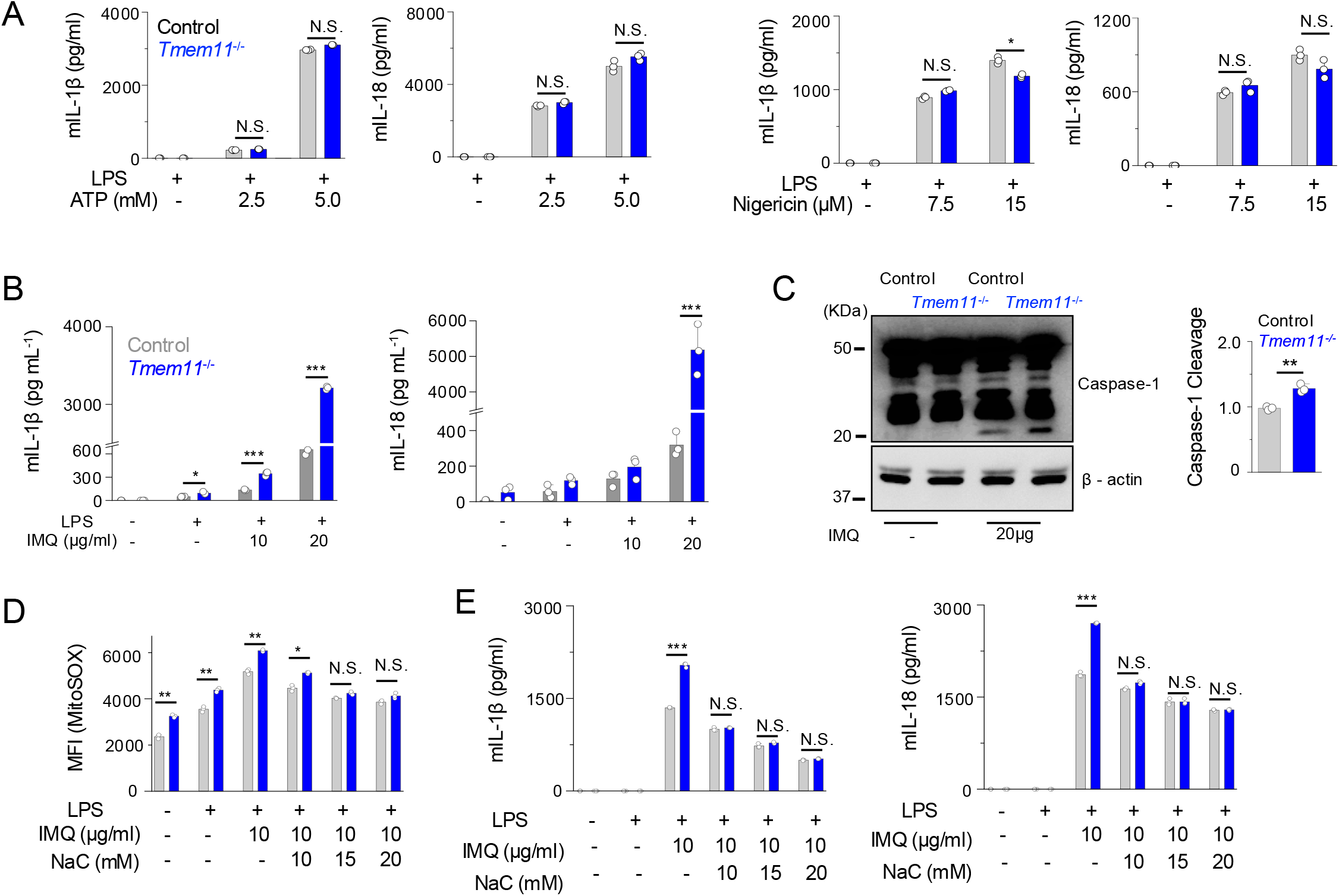
TMEM11 deficiency enhances imiquimod-induced activation of the NLRP3 inflammasome in a ROS-dependent manner. **A**, Secreted levels of IL-1β and IL-18 from LPS-primed control and *Tmem11*^-/-^ BMDMs stimulated with ATP (2.5 or 5 mM) for 30 min or nigericin (7.5 or 15 μM) for 45 mins. **B**, Secreted levels of IL-1β and IL-18 from LPS-primed control and *Tmem11*^-/-^ BMDMs stimulated with IMQ (10 or 20 μg/mL) for 90 mins. **C**, Representative immunoblot showing caspase-1 cleavage in LPS-primed control and *Tmem11*^-/-^ BMDMs upon IMQ stimulation. Normalized cleaved caspase-1 (p20) is shown as bar graph. **D**, Mitochondrial ROS measured by MitoSOX Red flow cytometry in control and *Tmem11*^-/-^ BMDMs under basal, LPS-primed, and IMQ-treated conditions. Cells were pre-treated with the indicated concentrations of N-acetylcysteine (NAC) for 30mins before IMQ stimulation (30mins). Mean fluorescence intensity (MFI) is shown. **E**, Effect of NAC on IMQ-induced IL-1β and IL-18 secretion in control and *Tmem11*^-/-^ BMDMs. Cells were pre-treated with the indicated concentrations of NAC for 30mins before IMQ stimulation. All experiments are representative of three independent experiments (unless otherwise indicated) and graphs show means ± s.e.m. from three independent experiments. *P < 0.05, **P < 0.005, ***P < 0.0005, N.S., not significant.

Activation of the NLRP3 inflammasome via the non-canonical pathway with intracellular LPS showed a mild increase in levels of secreted IL-1β from *Tmem11*^-/-^ macrophages (**Suppl. Fig. 1A**). In sharp contrast, stimulation of the K^+^ efflux-independent pathway of the NLRP3 inflammasome using IMQ, provoked an exacerbated, dose-dependent inflammatory response in *Tmem11*^-/-^ macrophages (**Fig. 3B**). Western blot analysis confirmed enhanced activation of the inflammasome, showing increase in cleaved active caspase-1 (p20) in lysates from IMQ-stimulated *Tmem11*^-/-^ macrophages. Interestingly, stimulation of the AIM2 inflammasome using poly dA:dT or Toll like receptors (TLRs) using RNA mimic poly I:C showed similar IL-1β secretion between control and *Tmem11*^-/-^ macrophages, suggesting specificity towards NLRP3 inflammasome activation (**Suppl. Fig. 1B**). Taken together, these data demonstrate that TMEM11-dependent cristae integrity selectively suppresses IMQ-induced inflammasome assembly without altering the cellular sensitivity to canonical K^+^ efflux-dependent stimuli.

**Supplemental Figure 1.**
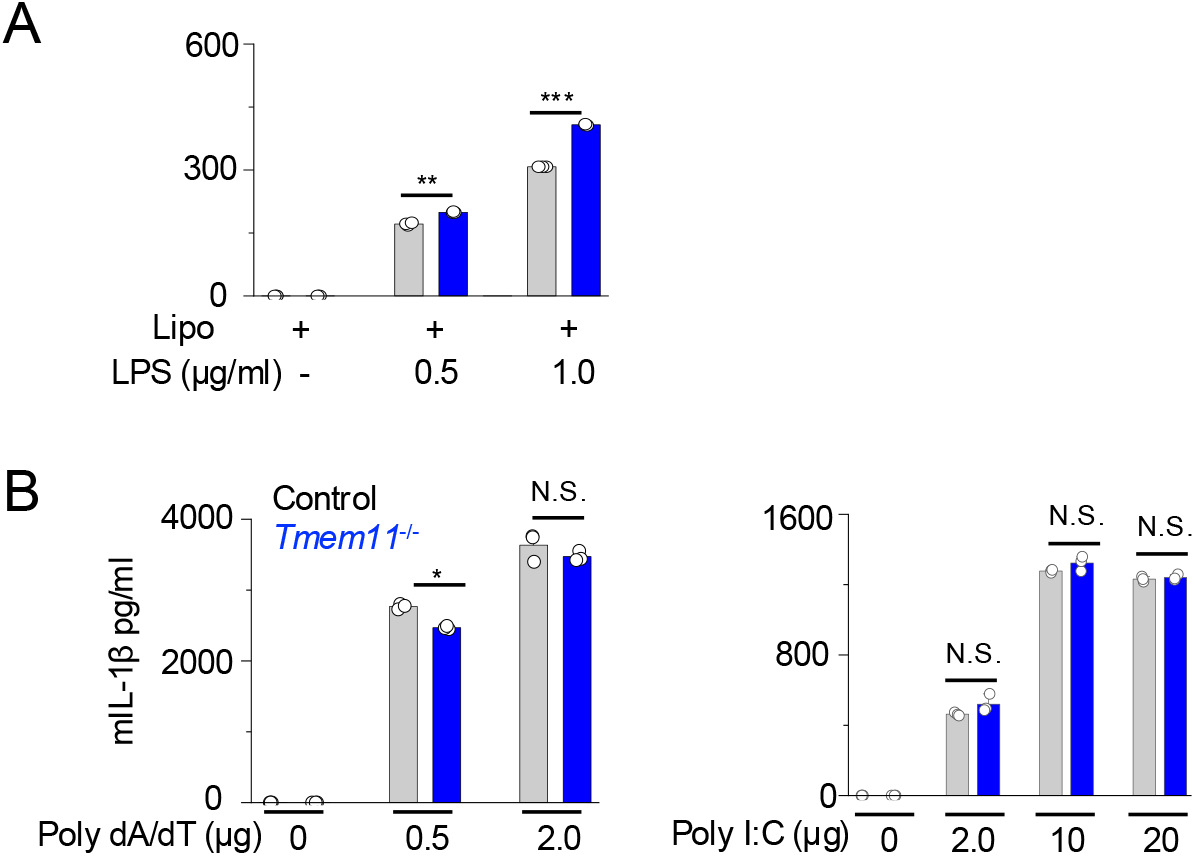
TMEM11 deficiency does not influence AIM2 inflammasome or poly I:C induced TLR responses. **A**, IL-1β secretion from control and *Tmem11*^-/-^ BMDMs following intracellular delivery of indicated amounts LPS using Lipofectamine 2000. **B**, IL-1β secretion from control and *Tmem11*^-/-^ BMDMs stimulated with indicated concentrations of poly(dA:dT) or poly(I:C) upon complexing with Lipofectamine 2000. Bar graphs in all the panels show means ± s.e.m. from 3 independent experiments; significance was determined by unpaired two-tailed *t*-test. *P < 0.05, **P < 0.005, ***P < 0.0005, N.S., not significant.

### Hyperactive inflammasome assembly in *Tmem11*^**-/-**^ macrophages is driven by exaggerated mtROS generation

Because our previous work demonstrated that *Tmem11* deficiency fundamentally elevates mtROS ^18^, we hypothesized that disrupted cristae architecture heightens electron leakage to amplify mtROS production upon IMQ-mediated complex I inhibition. In agreement with this hypothesis, flow cytometric profiling using the mitochondrial superoxide indicator MitoSOX Red demonstrated elevated mtROS in *Tmem11*^-/-^ macrophages (**Fig. 3D**). Under baseline and LPS-primed conditions, *Tmem11*^-/-^ macrophages showed higher MitoSOX mean fluorescence intensity (MFI) than controls. Addition of IMQ led to a significant increase in mtROS in *Tmem11*^-/-^ cells. To directly determine whether excessive mtROS was responsible for the heightened inflammasome activation, we treated macrophages with antioxidant N-acetylcysteine (NAC) in the presence of IMQ. As expected, NAC treatment suppressed IMQ-induced mtROS in *Tmem11*^-/-^ cells in a dose-dependent manner (**Fig. 3D**). Neutralizing mtROS with NAC completely abolished the hypersecretion of IL-1β and IL-18 in *Tmem11*^-/-^ macrophages to those observed in control cells (**Fig. 3E**).

These findings demonstrate that TMEM11-dependent cristae architecture limits IMQ-induced NLRP3 inflammasome activation by restraining mtROS production. In wild-type macrophages, organized cristae preserve stable ETC supercomplex assemblies, allowing efficient electron flux and controlled mtROS release during mild pharmacologic stress. When cristae junctions and sheets are structurally disrupted, pharmacologic inhibition of complex I triggers exaggerated electron leakage and superoxide generation, lowering the threshold for NLRP3 activation. Our results are in concurrence with the prior study that identified complex I as a target of IMQ for activation of the NLRP3 inflammasome^8^. In that study, treatment with glutathione peroxidase mimetics ebselen and pyrrolidine dithiocarbamate (PDTC) as well as NAC, reduced IMQ-induced activation of the NLRP3 inflammasome, suggesting an important role for mtROS in this event^8^. In contrast, Billingham et al. challenged the requirement for mtROS in IMQ- and CL097-induced NLRP3 inflammasome activation^12^. They proposed that while complex I inhibition by these compounds delivers an essential upstream trigger, it does so through an mtROS-independent mechanism that relies on intact mitochondrial electron transport to generate ATP, which is shuttled to the cytosol via phosphocreatine (PCr) and creatine kinase B (CKB) to license NLRP3 NACHT domain activity. However, this study used different ROS chelators including MitoTEMPO, S1QEL, and S3QEL to suppress mtROS levels. The exact efficacy of these ROS chelators in BMDMs was not directly demonstrated by concurrent measurement of mtROS. Our study combining a genetic model (*Tmem11*^-/-^) with the ROS chelator NAC clearly shows a requirement for ROS in IMQ-induced NLRP3 inflammasome activation, similar to observations made by Gross et al^8^.

Consistent with our previous findings demonstrating that TMEM11 deficiency establishes a genetically precise model of excessive mtROS generation without altering mitochondrial mass, biogenesis, or cell viability^18^, *Tmem11*^-/-^ macrophages maintain steady-state ATP levels under basal and LPS-primed states. Despite unchanged ATP pools, *Tmem11*^-/-^ macrophages generate marked bursts of mtROS and exhibit prominent hyperactivation of caspase-1 alongside excessive IL-1β and IL-18 secretion following IMQ exposure. Crucially, targeted neutralization of mtROS completely normalizes cytokine hypersecretion down to control levels, providing clear evidence that cristae-dependent mtROS production is both elevated and functionally necessary to drive xenobiotic-induced NLRP3 inflammasome hyperactivation.

## ACKNOWLEDGEMENTS

We thank Dani Torralba for help with maintenance of the TMEM11 knockout mouse colony and Dr. Linsey Stiles for use of Sea Horse extracellular flux analysis core facility. Flow cytometry was performed in the UCLA Jonsson Comprehensive Cancer Center (JCCC) and Center for AIDS Research Flow Cytometry Core Facility that is supported by National Institutes of Health awards P30 CA016042 and 5P30 AI028697, and by the JCCC, the UCLA AIDS Institute, the David Geffen School of Medicine at UCLA, the UCLA Chancellor’s Office, and the UCLA Vice Chancellor’s Office of Research. This work was supported by the National Institute of Health grants AI181072, AI201639 (Y.G.), and AI146352 (S.S).

## AUTHOR CONTRIBUTIONS

Y.G. and S.S. designed research and supervised the project. B.B.W. performed confocal analysis, FACS, most of the ELISA and immunoblotting analysis. S.S. performed electron microscopy, Super-Resolution STED Nanoscopy, and sea horse experiments in collaboration with C.B. M.D. helped with TMEM11 colony management, some ELISAs and seahorse data analysis. S.S., Y.G. and B.B.W. wrote the manuscript with input from all authors.

